# Notch is Required for Neural Progenitor Proliferation During Embryonic Eye Regrowth

**DOI:** 10.1101/2024.01.31.577915

**Authors:** Dylan J. Guerin, Belen Gutierrez, Baoyi Zhang, Kelly Ai-Sun Tseng

## Abstract

The ability of an organism to regrow tissues is regulated by various signaling pathways. One such pathway that has been studied widely both in the context of regeneration and development is the Notch signaling pathway. Notch signaling is required for development of the eye and regeneration of tissues in multiple organisms but it is unknown if Notch plays a role in the regulation of *Xenopus laevis* embryonic eye regrowth. We found that Notch1 is required for eye regrowth and regulates retinal progenitor cell proliferation. Chemical and molecular inhibition of Notch1 significantly decreased eye regrowth through reducing retinal progenitor cell proliferation without affecting retinal differentiation. Temporal inhibition studies showed that Notch function is required during the first day of regrowth. Interestingly, Notch1 loss-of-function phenocopied the effects of the inhibition of the proton pump, V-ATPase, where retinal proliferation but not differentiation was blocked during eye regrowth. Overexpression of a form of activated Notch1, the Notch intracellular domain (NICD) was sufficient to rescue loss of eye regrowth due to V-ATPase inhibition, suggesting that Notch acts downstream of V-ATPase. These findings highlight the importance of the Notch signaling pathway in eye regeneration and its role in inducing retinal progenitor cell proliferation in response to injury.

## INTRODUCTION

The ability of an organism to regrow lost or damaged tissues varies greatly among animals (Illingworth 1974; Michalopoulos and DeFrances, 1997; Slack et al., 2007; Tanaka and Reddien, 2011; Seifert et al., 2012; Frangoginannis, 2016; Tanaka, 2016; Tseng, 2017; Joven and Simon, 2018; Kha et al., 2018) In order to understand why some animals, or even some tissues within otherwise regenerative animals, lack this ability, there is a need to understand the molecular mechanisms that regulate this regrowth. An important model organism to study for regeneration is the African clawed frog, *Xenopus laevis. X. laevis* has long been studied as a regenerative model, valuable for their external fertilization and development, large clutch sizes, and relatively rapid development time, the speed of which can be manipulated by ambient temperature regulation (Gurdon and Hopwood, 2000; Sive, et al., 2000; Wlizla et al., 2018)

*Xenopus* displays age-dependent regeneration. Tadpoles can regrow a number of structures including the tail, limb, retina, and lens (Dent, 1962; Endo et al., 2000; Suzuki et al., 2006; Slack et al., 2007, Vergara and Del Rio-Tsonis; 2009; Mitogawa et al., 2018) with this ability generally decreasing in potency in some tissues as the animal ages (Slack et al., 2004; Beck et al., 2009; Kha and Tseng, 2018). Our previous work showed that *Xenopus* tailbud embryos regrew their eyes following surgical ablation of ∼85% of tissues (including the lens placode and most of the optic cup) at developmental stage (st.) 27 (Kha et al., 2018). The eye completes regrowth within 5 days, is functional, and contains the normal complement of cell types (Kha et al., 2018, Kha et al., 2019). The regrowth process requires cell proliferation and recapitulates retinogenesis. One advantage of this model is that eye regrowth in the embryo occurs concurrently with normal eye development at the uninjured contralateral side. In models where regrowth occurs post development, comparison of regrowth and development can be challenging due to the inherent differences between developing and mature tissues (Higgins and Anderson, 1931; Eguchi and Shingai, 1971; Yamada, 1977; Yoshii et al., 2006; Vergara and Del Rio-Tsonis, 2009). The embryonic eye regrowth model provides the opportunity for a more direct comparison between developmental and regrowth mechanisms in the same animal. This is important as developmental mechanisms are often co-opted into mechanisms regulating regrowth (Del Rio-Tsonis et al., 1997; Slack et al., 2004; Suzuki et al., 2007). Therefore, it is important to understand whether and how developmental mechanisms act as regulators of regrowth.

An important regulator of eye development is the Notch signaling pathway. Notch1 is a transmembrane receptor that upon binding to its ligand undergoes cleavage events resulting in the cleavage of the intracellular domain (NICD), which migrates to the nucleus and acts as a transcription factor to regulate downstream target genes (Bray, 2016). The Notch signaling pathway is a highly conserved, well-characterized developmental pathway that often determines if a cell population will proliferate or differentiate, and in some contexts, can maintain stem cell populations (Coffman et al., 1993; Henrique et al., 1997; Hitoshi et al., 2002; Borggrefe and Oswald, 2009, Reddy et al., 2010).

Notch signaling can also function as a regulator of stem cell proliferation. In the *Drosophila* wing disc, Notch works to upregulate proliferation, and overexpression of the Delta ligand was sufficient to increase proliferation in the wing disc (Baonza and Garcia-Bellido, 2000). In mouse, Notch promotes proliferation and maintains stemness in intestinal crypt base columnar stem cells. Reduction in Notch signaling reduced proliferation and expression of stem cell specific markers, and promoted differentiation (VanDussen et al., 2012). Similar behavior is found in bone marrow mesenchymal stem cells, where inhibition of Notch1 signaling resulted in reduced proliferation (He and Zou, 2019).

During *Xenopus* tadpole tail regeneration, Notch signaling is required for proper regrowth. Following tail amputation, treatment with MG132 (a proteosome inhibitor that blocks the cleavage of the Notch1 protein) resulted in healing of the tail stump without regeneration (Slack et al., 2004). Additionally, during the refractory period – when the tadpole temporarily loses its tail regenerative ability – activation of Notch signaling stimulated tail regeneration following amputation (Beck et al., 2003; Slack et al., 2004). During *Xenopus* eye development, active Notch serves to maintain a pool of multipotent retinal progenitor cells (RPCs) by regulating cell differentiation (Dorsky et al., 1995). An imbalance of Notch activity during development resulted in eye malformations (Dorsky et al., 1995, Furukawa et al., 2000). Given the roles of Notch in regulating stem cell populations and promoting appendage regeneration, it is likely that Notch acts to regulate RPC proliferation during *Xenopus* eye regrowth.

Here, we investigate the role of Notch signaling in *Xenopus* embryonic eye regrowth. Our study showed that loss of Notch1 function blocked eye regrowth and resulted in small eyes. Notch inhibition reduced RPC proliferation but retinal differentiation remained unaffected. We also determined that activation of Notch1 during regrowth is sufficient to rescue the regrowth-inhibited small eye phenotype caused by V-ATPase inhibition, demonstrating a link between Notch and V-ATPase signaling. Together, our results showed that Notch1 is required for eye regrowth.

## RESULTS

### Reduction of Notch following eye ablation inhibits regrowth

Successful eye regrowth requires the complex interaction of multiple cell signaling pathways including bioelectrical signaling and apoptosis (Kha et al., 2018; Kha and Tseng, 2018; Kha et al., 2023). Another candidate pathway is the Notch signaling pathway, which is required for development of the eye (reviewed in Blair 1999, Mills and Goldman, 2017, and Reichrath and Reichrath, 2020). In *Xenopus*, Notch1 is a neural stem cell marker. During development, it is expressed widely in the optic cup and inhibits retinal differentiation (Coffman et al., 1993; Zaghloul and Moody, 2007). However, the role of Notch1 in neural regrowth is unclear. Thus we investigated whether Notch1 is required for eye regrowth. First we sought to inhibit Notch signaling during eye regrowth. Previous studies of Notch function have successfully utilized the cysteine protease inhibitor MG132, as well as the γ-secretase inhibitor DAPT (Chapman et al., 2006, Slack et al., 2007, Xu et al., 2021) to inhibit Notch signaling by blocking the cleavage of the intracellular domain of the Notch protein, thereby inhibiting downstream activation. We first titrated each inhibitor to identify dosages that enabled normal development and used these concentrations (10 uM MG132 and 5 µM DAPT) for our experiments. Then we carried out eye regrowth assays to observe the effects of inhibitor exposure. Treatment with either chemical inhibitor following st. 27 ablation surgery caused a noticeable decrease in regrown eye size as compared to age-matched vehicle-treated regrowing controls (Fig. 1A-B). We used the Regrowth Index (RI, ranging from 0 to 300; described in Methods) to assess the overall quality of regrowth as judged by eye size and morphology. A 10 uM MG132 treatment resulted in 20.2% of fully regrown eyes (RI=172, n=114) as compared to DMSO (vehicle)-treated regrowing control which resulted in 63% of fully regrown eyes (RI=248, n=100, p < 0.01). Similarly, 5 uM DAPT treatment resulted in 47.9% of fully regrown eyes (RI=218, n=96) as compared to DMSO-treated regrowing control which resulted in 74.47 % of fully regrown eyes (RI=269, n=94, p < 0.01) (Fig. 1A-B). As chemical inhibitors can potentially have off target effects, these results were confirmed using molecular inhibition.

**Figure 1.**
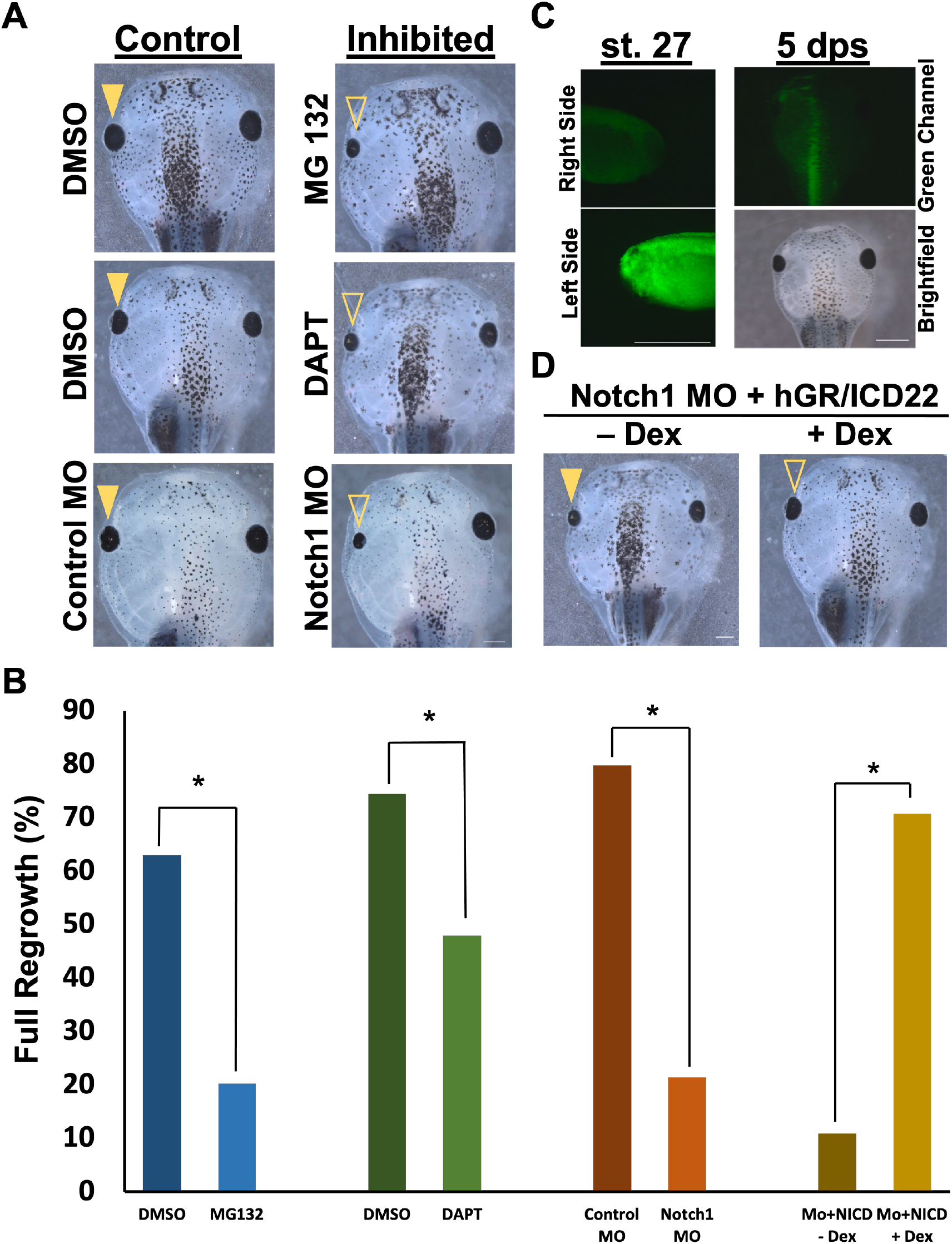
Reduced Notch impairs eye regrowth. (A) Comparison of 5 dps tadpoles treated with DMSO (control), Notch inhibitors (MG132 or DAPT), or injected with Notch morpholino (n>30 per condition). Images at 20.5X. Up= anterior. (B) Graph showing the percentage of the population achieving full regrowth under different conditions. *= p<0.01 (C) Morpholino expression. Cells containing morpholino showed green fluorescence from the fluorescein tagged oligonucleotide. Top left panel shows the right side of the embryo, right= anterior. Bottom left panel shows the left side of the same embryo, right= posterior. For both, up= dorsal. Images at 48X. Tadpole at 5 dps showing green fluorescence still present on the left side of the animal. Top right panel is green channel showing the morpholino fluorescein tag at 5 dps. Bottom right panel is the corresponding brightfield image. Up= anterior. Image at 25X. Scale bar= 500μm. (D) Comparison of 5 dps tadpoles expressing Notch1 morpholino and the Dex-inducible NICD in the left eye region. Left panel shows regrowth without the Dex, right panel shows regrowth with Dex. Closed arrowhead indicates control regrowth-inhibited eye. Open arrowhead indicates rescue of eye regrowth (n>30 per condition). Images at 20.5X. Up= anterior, down= posterior. (A, C, and D) scale bar= 500μm.

A verified morpholino against *Xenopus* Notch1 mRNA (Lopez et al., 2003) or a control morpholino was injected into the left dorsal blastomere of four-cell embryos at a concentration that does not affect embryogenesis. The morpholinos were tagged with fluorescein, allowing for the selection of embryos with eye regions that contained high levels of either the control or Notch1 morpholino (Fig. 1C). At st. 27, eye ablation surgeries were performed on these embryos and regrowth was assayed at 5 dps. Consistent with the chemical inhibition results, embryos with the Notch1 morpholino had significantly reduced eye regrowth (21.4% full regrowth, RI=165, n=109) as compared to those injected with the control morpholino (79.8% full regrowth, RI=276, n=112, p < 0.01) (Fig. 1A-B). To confirm that the morpholino effect was due to Notch1 knockdown, we tested if ectopic expression of Notch1 could restore eye regrowth. The Notch Intracellular Domain (NICD) is an activated form of Notch and an established tool to activate Notch signaling in *Xenopus* (Coffman et al., 1993). Dexamethasone-inducible NICD mRNA was co-injected with the Notch1 morpholino during the four-cell stage. To induce NICD activation, 10 µM dexamethasone (Dex) was added after eye ablation surgery. Dexamethasone treatment resulted in 59.1% full eye regrowth (RI=248, n=22) as compared to 9.5% full regrowth in the uninduced control (RI=144, n=21; p < 0.01) (Fig. 1B). Thus NICD activation was sufficient to rescue Notch1 morpholino-inhibited regrowth (Fig. 1D). This result indicated that it was the reduction in Notch1 which caused the inhibition of regrowth. Together, our data demonstrated that Notch is required for successful regrowth of the eye.

### Notch is required during the first day of regrowth

Notch signaling is often a regulator of proliferation (Baonza and Garcia-Bellido, 2000; Pallavi et al., 2012; VanDussen et al., 2012). Our previous work showed that approximately 87% increase in eye size during regrowth occurred during the first two days (Kha et al., 2018). We hypothesized that the requirements for Notch function during eye regrowth is during the early time period. To identify the temporal requirement for Notch, the duration of exposure to 10 µM MG132 was varied during the regrowth period. Our data indicated that embryos with reduced Notch function during the first 24 hours post-surgery resulted in similar inhibition (RI=191, n=99) with 25.3% of full regrown eyes as compared to embryos inhibited for the entire five-day period with 24.7% of full regrown eyes (RI=191, n=93, p > 0.05) (Fig. 2). If the key requirement for Notch activity is during the first day of regrowth, then its inhibition after 1 dps should not affect regrowth. Consistent with this prediction, embryos treated with MG132 from 1 dps through the end of the five-day assay showed no appreciable inhibition of regrowth with 78.8% of full regrown eyes, a level that was comparable to DMSO-treated controls (RI=250, n=92; p < 0.01 compared to either 0-5 dps or 0-1 dps treatment) (Fig. 2B). Together, our data showed that the first 24 hours is the critical period where Notch function is required to drive eye regrowth.

**Figure 2.**
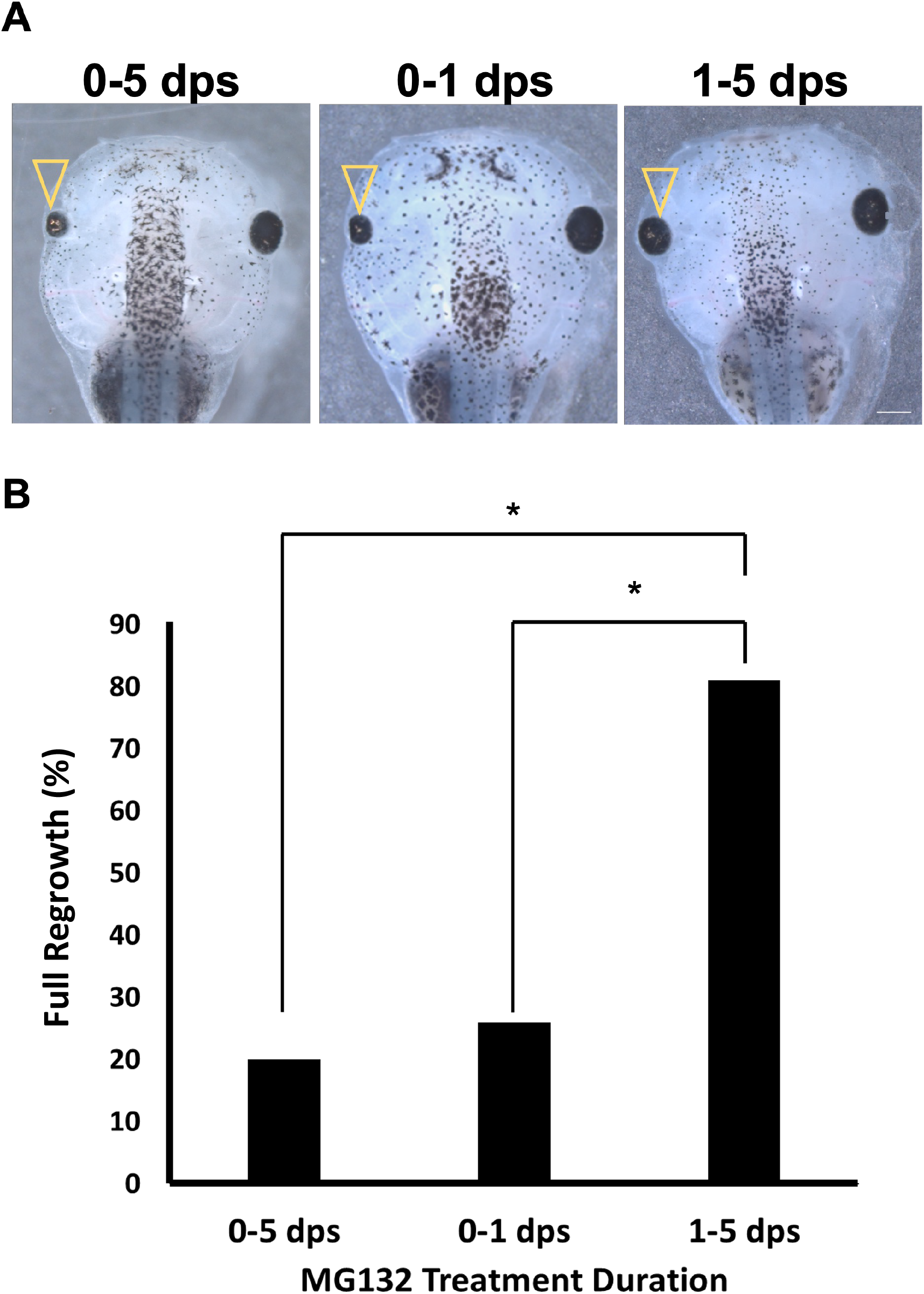
Notch is required during the first day of regrowth. (A) Comparison of regrown eyes at 5 dps following treatment of MG132 for variable durations from time of surgery (n>30 per condition). Open arrowheads indicate regrown eye. Up= anterior, images at 20.5X. Scale bar= 500μm. (B) Graphical representation of percent of the population achieving full regrowth at 5 dps, * = p<0.01.

### Disruption of Notch does not disrupt differentiation

In order for eye regrowth to occur fully, both cell proliferation and proper retinal differentiation are needed. Previously, we showed that the regrown eye contained the expected complement of retinal cell types including rod and cone photoreceptors, ganglion cells, retinal pigmented epithelium cells, and Müller glia, with the same structure as a normally developing eye (Kha et al., 2018). Defective regrowth could result from disruptions in retinal cell proliferation or differentiation, or both. To characterize the regrowth defects resulting from Notch inhibition, we first examined the effects on retinal differentiation. It is possible that reducing Notch function during regrowth disrupted retinal differentiation, leading to small eyes. In order to test for this, we examined retinal differentiation using known antibody markers. Embryos were fixed at 3 dps following either treatment with MG132 or injection with Notch1 morpholino, as well as control regrowing eyes. Eye sections were obtained and stained with anti-Islet1 antibody as a marker of ganglion cells, or anti-Rhodopsin antibody as a marker of rod photoreceptor cells (Kha et al., 2019). Similar retinal patterns were observed between the Notch-inhibited non-regrowing small eyes and control regrowing eyes in terms of overall morphology and positioning of ganglion and rod photoreceptor cells relative to their eye sizes (Fig. 3). Islet1 signal was present within the ganglion cell layer and inner nuclear layer in both the control regrowing and Notch-inhibited non-regrowing eyes (Fig. 3A, n > 5 for each condition). As expected, the rhodopsin signal spanned the posterior periphery of the eye, in the outer nuclear layer (Fig. 3B, n > 5 for each condition). These observations are consistent with previous work in *Xenopus* showing that abnormal small eyes can contain largely normal retinal layers (Harris and Hartenstein, 1991; Kha et al., 2019). Additionally, regrowing eyes showed a delay of differentiation as compared to normal developing eyes (Kha et al., 2019). Here, we find that non-regrowing eyes also showed delayed differentiation, same as the regrowing eyes. Together, these results indicated that perturbation of Notch function during eye regrowth does not disrupt retinal differentiation.

**Figure 3.**
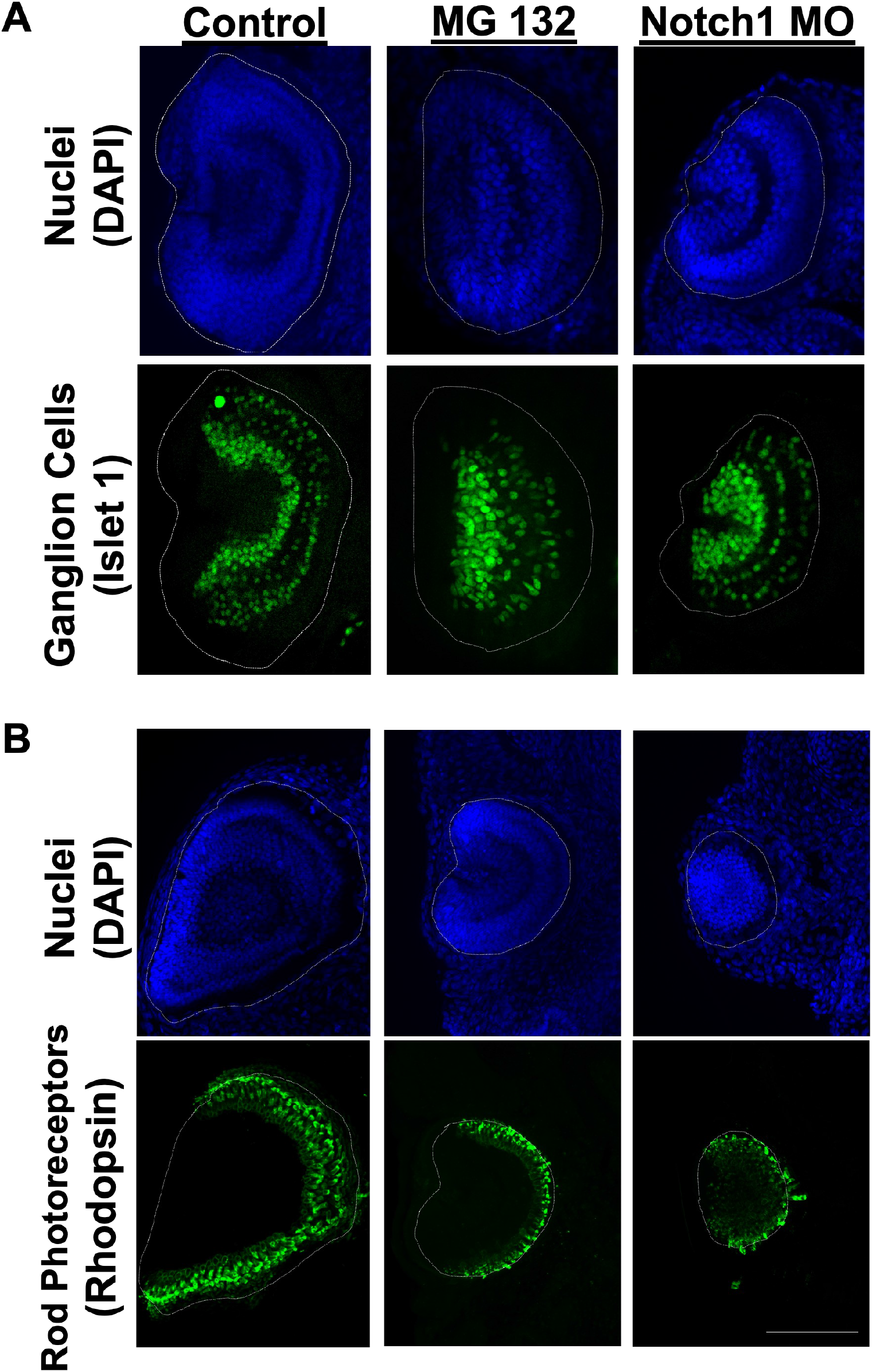
Reduction in Notch during eye regrowth does not impair differentiation. (A) Transverse eye sections of regrowing and inhibited eyes at 3 dps. Top panels indicate nuclei. Bottom panels indicate ganglion and inner nuclear layer cells. (B) Transverse eye sections of regrowing eyes at 3 dps. Dotted lines outline the eye region. Top panels indicate nuclei. Bottom panels indicate rod photoreceptor cells. All eyes were sectioned medially and facing left. Images at 20X. Scale bar= 100μm. n>5 per condition.Up= dorsal, Left= anterior.

### Inhibition of Notch function downregulates retinal proliferation

In order for the eye to regrow successfully, cellular proliferation must take place to replace the tissues that were lost. There is a significant increase in the number of proliferating cells in the regrowing eye during 0-1 dps relative to normally developing eyes (Kha et al., 2018). This increase is within the same window in which Notch is required for eye regrowth. Notch is known to play a role in regulating proliferation during eye development (Go et al., 1998). As Notch does not appear to regulate retinal differentiation during regrowth, we examined whether Notch regulates cell proliferation during regrowth.

Embryos injected in the left dorsal blastomere at the four-cell stage with Notch1 morpholino or treated with MG132 following surgery were fixed at 1 dps, sectioned, and stained with anti-phospho-Histone H3 (H3P) antibody, an established marker of mitosis (Adams et al., 2007; Kha et al., 2023; Miller et al., 2023). The number of H3P-positive cells was counted and then normalized to the area of the eye to account for the decreased size of the regrowth-inhibited eyes. Consistent with morphological observations (Fig. 3A and C), embryos with chemical (n= 13) or molecular (n=9) Notch inhibition showed either 56.7% or 45.8% reduction in the number of mitoses relative to their respective control regrowing eyes (n=10 and n=8; p < 0.05) (Fig. 4B and D). These data indicated that reduction in Notch during the regrowth period led to a reduction in RPC proliferation within the eye during regrowth.

**Figure 4.**
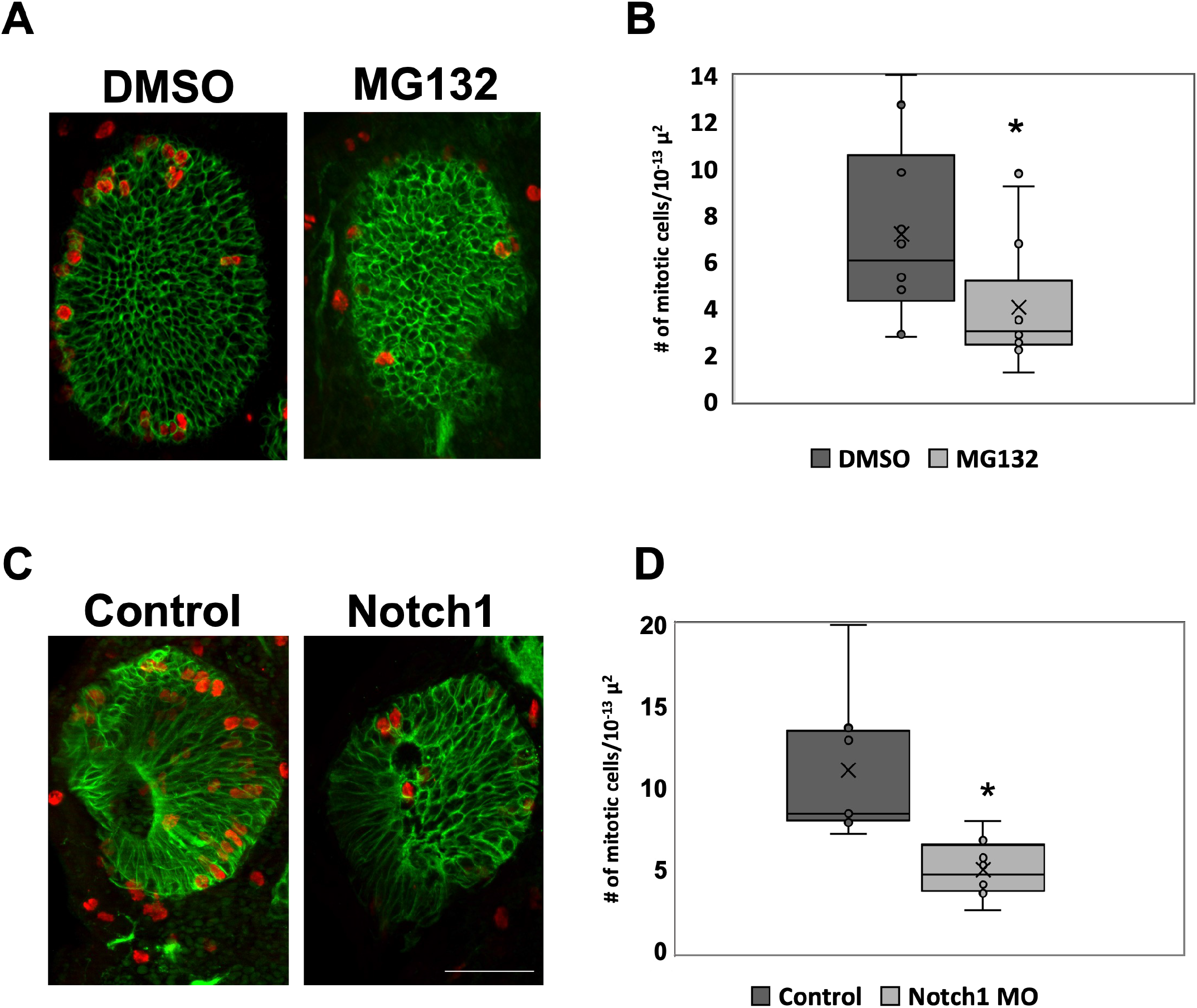
Reduction in Notch during eye regrowth reduces proliferation. (A) Transverse eye sections at 1 dps showing mitosis cells in DMSO or MG132 treated eyes (n>5 per condition). Red= phosphorylated Histone 3, Green= Xen1. Images at 20X. (B). Box and whisker plot comparing number of H3P-positive cells normalized to area. (C) Transverse eye sections at 1 dps comparing proliferating cells in control morpholino and Notch1 morpholino injected eyes (n>5 per condition). Red= phosphorylated Histone 3, Green= Xen1. Images at 20X. Scale bar= 100μm. (D) Box and whisker plot comparing number of H3P-positive cells normalized to area. *= p<0.01. Up= dorsal, Left= anterior.

### Notch1 overexpression restores eye regrowth during V-ATPase inhibition

Although much attention has focused on the roles of well-characterized developmental signaling pathways in regrowth, there are other key mechanisms that also determine regenerative success. Our lab recently showed that the proton pump V-ATPase is required for eye regrowth but did not appear to have a role in eye development (Kha et al., 2023). In mouse, blockage of V-ATPase activity reduced Notch signaling leading to reduced proliferation of neural stem cells (Lange et al., 2011). We observed that the block of eye regrowth caused by Notch inhibition gave the same phenotypes as the effects of V-ATPase inhibition: small eyes and reduced cell proliferation with normal retinal patterning. As such, we asked whether Notch and V-ATPase can interact to regulate eye regrowth.

We hypothesized that V-ATPase acts upstream of Notch signaling during eye regrowth. Thus, we tested whether it was possible to rescue V-ATPase inhibition of regrowth through ectopic activation of Notch signaling. We co-injected the following: GFP mRNA along with mRNA for Dex-inducible NICD into the left dorsal blastomere at the four-cell stage; and later selected for those embryos expressing GFP in the left eye region (Fig. 5A). To inhibit V-ATPase activity, the highly specific inhibitor, Concanamycin A (Huss et al., 2002; Adams et al., 2006; Kha et al., 2023) was used. 20 nM Concanamycin A successfully blocked eye regrowth without affecting development (Kha et al., 2023). Following eye ablation, embryos were treated with Concanamycin A and 10 µM dexamethasone, allowing for the ectopic activation of Notch signaling in the eye region after surgery. Controls were treated with Concanamycin A only. Regrowth was assessed at 5 dps (Fig 5B-C). Control non-induced embryos treated with Concanamycin A resulted in 22.6% of full regrown eyes with a low RI of 131 (n=110). In contrast, embryos expressing NICD in the presence of Concanamycin A resulted in 75.5% of full regrown eyes with an RI of 260 (n=106, p < 0.01), representing a comparable quality of regrowth as untreated regrown eyes. Our data showed a significant rescue of V-ATPase inhibition in the embryos overexpressing NICD as compared to the control. This result showed that activation of Notch signaling is sufficient to rescue regrowth following V-ATPase inhibition. Moreover, Notch acts downstream of V-ATPase to regulate regrowth.

**Figure 5.**
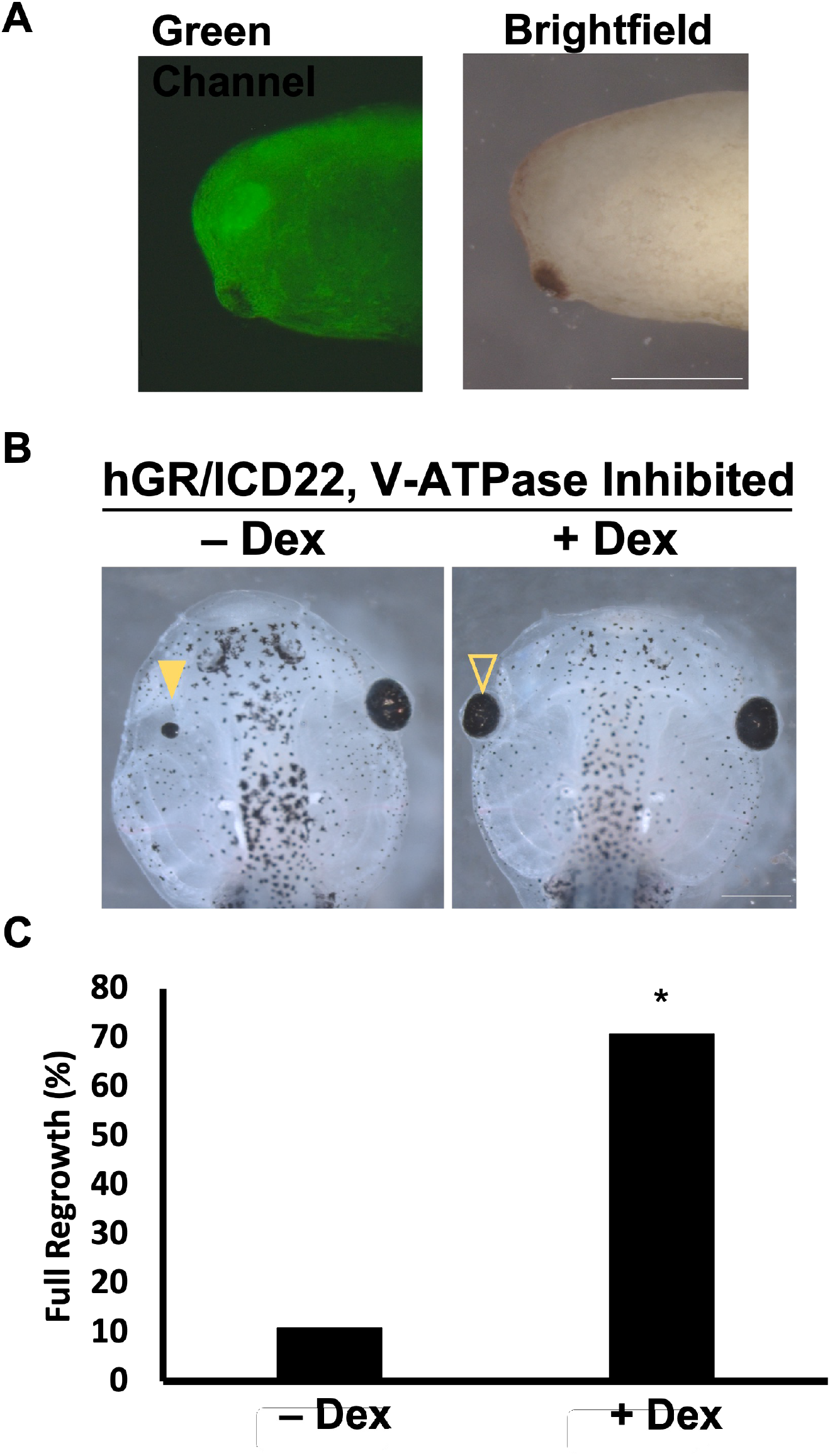
Notch1 overexpression during regrowth rescues V-ATPase inhibition. (A) Notch1 RNA is present in the eye at time of surgery. Left side of a st. 27 embryo injected at st. 3 with GR-NICD RNA and GFP RNA. Right= posterior, left= anterior, up= dorsal, down= ventral. Images at 48X. (B) Comparison of tadpoles at 5 dps of concanamycin treatment, injected with Dex-inducible NICD at st. 3 and treated with or without inducer immediately following ablation. Closed arrowhead indicates control regrowth-inhibited eye. Open arrowhead indicates rescued regrown eye though NICD overexpression. n>30 per condition. Images at 25X. Scale bar= 500μm. (C) Graphs showing percent of the population achieving full regrowth at 5 dps with or without NICD activation. *= p<0.01. Up= dorsal, Left= anterior.

## DISCUSSION

In this study, we show that Notch1 is a required component of successful eye regrowth in *Xenopus laevis*. This finding is consistent with previous studies linking well-known eye developmental pathways such as FGF, Pax6, retinoic acid, Wnt, and JAK/STAT, as necessary for successful retinal regeneration (Kaneko et al., 1999; Osakada et al., 2007; Spence et al., 2008; Hochmann et al., 2012; Todd et al., 2016; Tseng, 2017; Todd et al., 2018; Kha et al., 2019; Gao et al., 2021). During *Xenopus* eye formation, Notch promotes RPC proliferation by inhibiting differentiation (Dorsky et al., 1995). Notch is also active in the ciliary margin zone (a self-renewing proliferative region located at the periphery) of the mature tadpole retina and acts to maintain retinal progenitor cells (Perron et al., 1998). We determined that Notch signaling increased retinal progenitor proliferation during regrowth. Although Notch can display pleiotropic effects, its function was not needed for retinal differentiation during regrowth. This is consistent with the observation that Notch is required during the first day of eye regrowth but not later on, when delayed retinogenesis becomes active.

In *Xenopus*, retinal differentiation starts at st. 24 and is completed by st. 42, over a period of two days. At st. 27, RPC cell division time is 8.6 hrs and it increases to 56 hrs by st. 37/38, when most cells have exited the cell cycle (Rapaport, 2006). In other types of neural stem cells, inhibition of Notch signaling has been shown to lengthen the cell cycle time (Borghese et al., 2010; Alhashem et al., 2022). These results suggest that reducing Notch during regrowth may lead to a lengthening of RPC doubling time with the consequence of a smaller RPC pool causing a failure to restore the eye to the appropriate size. In this case, the likely role of Notch in regrowth would be to maintain the short RPC doubling time (as in st. 27) to allow for the restoration of the retinal progenitor population after eye ablation.

Multiple studies suggest a dynamic role for Notch signaling in retinal regeneration. After zebrafish retinal injury, the Müller glia responds by asymmetrically dividing to provide a neural progenitor cell population capable of regeneration (Fausett and Goldman 2006; Bernardos et al., 2007, Fimbel et al., 2007, Thummel et al., 2008). Notch signaling is upregulated during regeneration but normally acts to maintain quiescence in adult Müller glia populations by downregulating proliferation when there is no damage (Wan et al., 2012). Chick and rodent retinas undergo limited retinal regeneration. Inhibition of Notch signaling reduced progenitor cell proliferation (Hayes et al., 2007, Karl et al., 2008; Del Debbio et al., 2010). However, continued Notch signaling subsequently prevented neuronal differentiation in the chick retina. Nevertheless, Notch signaling is consistently supportive of increasing progenitor pools during retinal regeneration as is the case for *Xenopus* eye regrowth.

The mechanisms that regulate eye regrowth are beginning to be identified (Kha et al., 2018; Kha et al., 2019; Kha et al., 2023). Here, we found that ectopic expression of NICD rescued eye regrowth failure resulting from inhibition of V-ATPase. Other studies have observed similar interactions between V-ATPase and Notch. V-ATPase is expressed on cellular membranes as an essential H^+^pump. In *Drosophila*, reduction in V-ATPase activity caused disruptions in endocytic acidity, leading to defective trafficking and processing of the Notch protein (Yan et al., 2009). V-ATPase inhibition in mouse led to a reduction of Notch signaling which could be rescued by the expression of NICD but not a plasma membrane-bound form of activated Notch, suggesting that V-ATPase acted upstream of γ-secretase-dependent cleavage of the NICD (Lange et al., 2011). The regulation of Notch processing by V-ATPase could be the mechanism that is used during eye regrowth. However, other findings suggest alternative mechanisms. The ectopic expression of a yeast plasma membrane H^+^ pump, PMA, was sufficient to rescue tadpole tail and eye regrowth failures induced by V-ATPase inhibition (Adams et al., 2007; Kha 2023). As PMA is located on the cell surface, this finding suggests that it is the plasma membrane functions of V-ATPase that is essential for eye regrowth rather than its vesicular membrane roles. During tadpole tail regeneration, Notch1 RNA expression in the regeneration bud is absent when bioelectrical signaling was inhibited (Tseng et al., 2010). Additionally, transcriptional profiling of *Drosophila* neuroblasts suggested that Notch and V-ATPase also interacted in a regulatory loop (Wissel et al., 2018). Further studies will be needed to explore the specific interactions between Notch and V-ATPase in promoting eye regrowth.

Many developmental signaling pathways play an essential role in regrowth of tissues after injury. However, the specific interactions may not be the same during tissue restoration. As in development, we found that Notch promotes neural proliferation during eye regrowth (Dorsky et al., 1995; Wall et al., 2009; Schouwey et al., 2011). We also uncovered an eye regrowth pathway where Notch acts downstream of V-ATPase. Although Notch plays a key role in *Xenopus* eye development, there is no known role for V-ATPase in the same process. Here, activation of V-ATPase appears to be a regrowth-specific signal. Therefore, it would be informative to individually determine which of the common developmental pathways are active within eye regrowth and how they interact with regeneration-specific mechanisms. Furthermore, this knowledge may help to inform potential strategies for ocular repair.

## EXPERIMENTAL PROCEDURES

### Embryo culture and surgery

This study was carried out in accordance with the recommendations of the University of Nevada, Las Vegas Institutional Animal Care and Use Committee (IACUC). Embryos were obtained via *in vitro* fertilization and raised in 0.1X Marc’s Modified Ringer (1 mM MgSO4, 2.0 mM KCl, 2 mM CaCl2, 0.1 M NaCl, 5 mM HEPES, pH 7.8) (Sive et al., 2000). Eye ablation surgery was performed as described in Kha et al., 2020. Following surgery, embryos were cultured at 22^0^C.

### Assessment of Eye Regrowth

The regrowth quality of eyes treated with chemical or molecular inhibitors after surgery were compared to age-matched regrown eyes from the same batch of embryos by measuring the percent of the population achieving full regrowth and calculating the Regrowth Index (RI), a quantitative measurement where the percentage of embryos achieving each category of regrowth is assigned a numerical value, with values for the group ranging from 0-300, with 0 indicating all embryos failed to regrow eyes and 300 being all embryos fully regrew eyes, as described in (Kha et al., 2020).

### Chemical Treatments, and Morpholino and RNA Injections

Inhibitors were dissolved in DMSO. For knockdown of Notch, embryos were cultured in medium containing DAPT (Cayman Chemical, Ann Arbor Michigan, CAS number 208255-80-5) or MG132 (Cayman Chemical, Ann Arbor Michigan, CAS number 1211877-36-9) immediately following surgery and cultured in the medium for five days. Control embryos were immersed in medium containing an equivalent concentration of DMSO. To determine the temporal requirement for Notch, embryos were cultured in MG132 with varying time periods. As needed, embryos were removed from chemical-containing media, washed in 0.1X MMR, and transferred to 0.1x MMR, or taken from 0.1x MMR at the 1 dps time point and transferred into media containing MG132 for the remainder of the regrowth period (4 days). At 5 dps, embryos were washed with fresh media, anesthetized, and regrowth was assayed.

For morpholino injections, the following published morpholinos were purchased from Gene Tools LLC (Philomath, Oregon): Notch1 5’-GCACAGCCAGCCCTATCCGATCCAT-3’ (Lopez et al., 2003) and the non-specific standard control oligomer: 5’-CCTCTTACCTCAGTTACAATTTATA-3’. Each morpholino contained a 3’ fluorescein addition. 2.5 ng of morpholinos were injected into the left dorsal blastomere at the four-cell stage. Embryos with fluorescent signal in the eye region at st. 27 were selected for further analysis. hGR/ICD22 and GFP were transcribed *in vitro* from linearized plasmid constructs using the mMESSAGE Transcription Kit (Thermofisher). For injections, GFP and 0.25 ng hGR/ICD22 (Coffmanet al., 1993) mRNA were co-injected into the left dorsal blastomere at the four-cell stage. Embryos with fluorescent signal in the eye region at st. 27 were selected for further analysis. Dexamethasone was added to media as an inducer at a final concentration of 10 μM.

## Embryo Sectioning and Immunofluorescence Microscopy

For agarose embedding and sectioning, animals were fixed overnight at 4°C in MEMFA (100 mM MOPS (pH 7.4), 2 mM EGTA, 1 mM MgSO4, 3.7% (v/v) formaldehyde) (Sive et al., 2000) and dehydrated in methanol. After rehydration, embryos and tadpoles were embedded in 4–6% low-melt agarose and sectioned into 60 µm slices using a Leica vt1000s vibratome. Sections were stained with primary antibodies including: Xen1 (pan-neural antibody, clone 3B1, 1:50 dilution, Developmental Studies Hybridoma Bank, RRID: AB_531871), anti-Islet1 (retinal ganglion cells and inner nuclear cell layer, clone 40.2D6, 1:200 dilution, Developmental Studies Hybridoma Bank, RRID: AB_528315), anti-Rhodopsin (rod photoreceptor cells, clone 4D2, 1:200 dilution, EMD Millipore, RRID: AB_10807045), and anti-phospho Histone H3 (mitosis marker, 1:500 dilution, EMD Millipore, RRID:AB_310177). Alexa fluor conjugated secondary antibodies were used at 1:1000 dilution (ThermoFisher Scientific). n > 5 was used for each antibody.

## Microscopy

A Nikon A1R confocal laser scanning microscope (UNLV Confocal and Biological Imaging Core) was used to image Islet1 immunostained tissue sections. All other immunostained tissue sections were visualized via Zeiss Axio Upright Imager M2 microscope with a Hamamatsu ORCA flash 4.0 monochromatic digital CMOS camera. Images of whole animals were obtained using a ZEISS SteREO Discovery V20 microscope with an AxioCam MRc camera. ZEN Image Analysis software and/or the open-source FIJI imaging software (Schindelin et al., 2012) were used to analyze and/or process all acquired images.

## Statistical Analysis

To compare eye regrowth, raw data from scoring was used. Comparison of two treatments was analyzed with Mann-Whitney U test for ordinal data with tied ranks, using normal approximation for large sample sizes. All other experiments were analyzed using a Student’s t-test.

## ACKNOWLEDGEMENTS

We thank the members of the Tseng lab for their help and Christopher Kintner and Richard Harland for the Notch overexpression construct. This study was supported by a grant from the National Institutes of Health (GM146672) to KT, and research fellowships from the Nevada NASA Space Grant Consortium (80NSSC20M0043) to DG, and National Science Foundation (1301726 and 1757316 to BG, and 2148788 to BZ). Confocal imaging was performed at the UNLV Confocal and Biological Imaging Core, with assistance of Sophie Choe. Some antibodies used in this study were obtained from the Developmental Studies Hybridoma Bank, a resource created by the NICHD of the NIH and maintained at The University of Iowa, Department of Biology, Iowa City, IA.

